# Age-dependent variation of aedeagal morphology in *Agabus uliginosus* and the status of *A. lotti* (Coleoptera, Dytiscidae)

**DOI:** 10.1101/2024.05.07.592935

**Authors:** Konrad Dettner, Zsolt Kovács, Tomasz Rewicz, Zoltan Csabai

## Abstract

The taxonomic status of *Agabus lotti* within the *Agabus uliginosus* species group has been a subject of debate due to morphological similarities and lack of molecular data. In this study, we conducted a comprehensive morphological and molecular analysis of specimens from Central Europe, focusing on the Hungarian population. Morphological comparisons of genital structures revealed age-dependent variations, suggesting a gradual transition from *A. lotti* to *A. uliginosus*. Molecular analysis of COI sequences further supported this hypothesis, showing minimal genetic differences among most specimens, with only one individual exhibiting distinctiveness. Therefore, *A. lotti* should be regarded as a junior synonym of *A. uliginosus*. Our findings also highlight the need for additional multi-marker studies covering a broader geographic range and including both molecular and morphological approaches to elucidate the taxonomic and phylogenetic relationships within this species group. The inclusion of Hungarian samples notably enriched the diversity of haplotypes, emphasizing the importance of expanding sampling efforts in future research.

## Introduction

The uliginosus-group, within the genus Agabus (Dytiscidae, Agabinae), as delineated in its present form by Larson (1989) and Nilsson (2000), and currently summarized by Nilsson & Hájek (2024), comprises seven species: *A. amnicola* (J. Sahlberg, 1880) with a Holarctic distribution, *A. jacobsoni* Zaitzev, 1905, *A. lotti* Turner, Toledo, et Mazzoldi, 2015, *A. uliginosus* (Linnaeus, 1761), *A. uralensis* Nilsson et Petrov, 2006, and *A. vereschaginae* Angus, 1984 with a distribution in the Palaearctic region, while *A. margareti* Larson, 1975 (=*A. margaretae* Larson, Alarie & Roughley, 2000 and Turner, Toledo & Mazzoldi, 2015, as unjustified emendations of *margareti*, see also Fery & Bouzid, 2016) is a Nearctic species. Previously, Larson (1989), Nilsson & Holmen (1995), and Nilsson & Toledo (1999) grouped *A. amnicola*, *A. jacobsoni*, *A. uliginosus*, *A. vereschaginae*, *A. margareti*, and *A. falli* (Zimmermann, 1934) within the uliginosus group; however, Nilsson (2000) later excluded *A. falli* from this group and transferred it into its own group. Subsequently, Nilsson & Petrov (2005) distinguished the Asian *A. uralensis* from A. *uliginosus* based on male genitalia, thereby elucidating the distribution of *A. uliginosus* in the western Palaearctic. The description of *A. lotti* as the sixth Palaearctic species within the uliginosus group took place in 2015 (Turner, Toledo & Mazzoldi, 2015). This species, endemic to Central Europe, can be readily distinguished from *A. uliginosus* by its small phallobase and thin bases of parameres, in contrast to the considerably enlarged phallobase and thickened, more sclerotized bases of adhering parameres in *A. uliginosus*. As described by Turner et al., (2015), „penis in lateral view basally sinuate expanding medially, followed by a sinuate contraction toward the narrowing apex; the external margin progresses evenly in a gradual inward curve until the three quarters to the apex where a straight section completes the apex”. Among the external characters, the authors highlight the more elongated body shape of *A. uliginosus*, compared to the laterally more expanded body shape of *A. lotti*.

According to Lawrence et al., (2010), a coleopteran aedeagus comprises the phallobase, parameres, and penis. In the trilobate type aedeagi, the parameres are paired structures articulated with the posterior end of the phallobase, which is the basal portion of the aedeagus. Lawrence et al., (2010) notes that in many Coleoptera (including Adephaga), the phallobase appears to be absent, having become membranous or fused to the parameres. Additionally, they define the tegmenite as a slender sclerite articulated with sternite IX or with the connecting membrane between sternite IX and the phallobase. These structures were consequently illustrated by Franciscolo (1979) for European species. Miller & Bergsten (2016) described various types of male genitalia within Dytiscidae and denoted that the median lobe may be either bilaterally symmetrical (Dytiscinae, most Hydroporinae) or asymmetrical and variously twisted.

In insects, particularly adephagous beetles, it is well-established that internal reproductive organs develop gradually and significantly increase in size with age, as progressing from immature to teneral adults and eventually to fully sclerotized, old individuals. Various age classes were consequently described for both females and males (Dettner, Hübner & Classen, 1986). While degeneration of internal genitalia in elder beetles typically occurs post-reproduction, instances of genital reactivation have been observed in beetles living for extended periods, sometimes several years (Blunck, 1912). Fresh weights, though subject to fluctuations due to varying food intake, can serve as an approximate indicator of age, owing to the gradual increase in biomass of ectadenies (=accessory glands), testes, and ducti ejaculatori in males, and ovaries, eggs, oviducts, and vagina in females (Dettner, Hübner & Classen, 1986).

DNA barcoding offers an effective method for species-level identification and mapping genetic diversity, enabling the detection of atypical specimens for thorough taxonomic analyses and as such, give much potential to reveal new species or to review the taxonomic status of previously described taxa (Hajibabaei et al., 2007). Among the genetic markers used in DNA barcoding, mitochondrial cytochrome c oxidase subunit I (COI) has proven to be particularly effective for species identification across the animal kingdom, with its divergences facilitating discrimination even among closely related species in nearly all animal phyla (Hebert, Ratnasingham & deWaard, 2003; Hebert et al., 2003).

Despite the potential of DNA barcoding, COI-based DNA reference databases remain incomplete (e.g., Weigand et al., 2019), including for adephagous aquatic beetles (Csabai et al., 2023). Currently, publicly available sequences exist for only two of the seven species within the *uliginosus*-group in databases such as BOLD (Ratnasingham & Hebert, 2007) and GenBank (Benson et al., 2013). There are 17 public/published sequences from Germany and Finland for *A. uliginosus*, all of which are classified within a single BIN (Barcode Index Number, Ratnasingham & Hebert, 2013), with an average distance of 0.38% and a maximum distance of 1.52%. Additionally, a single sequence from Canada is available for *A. margareti*, forming a separate BIN (BOLD:ACA6100), with a 6.9% distance from the nearest neighbour, the *A. uliginosus* BIN (BOLD:AAY8849). Venables (2016), in her PhD thesis on the molecular systematics of the Agabini, included *Agabus uliginosus* and *A. lotti* in some phylogenetic analyses, albeit with only single specimens analysed. These two species, along with *A. lineatus* (*lineatus*-group), represented a monophylum, with *A. lotti* and *A. uliginosus* positioned very closely.

Given the morphological similarity and genetic proximity of these species, it is more appropriate to reach a consensus based on various methods rather than relying solely on a single approach. Therefore, in this study, alongside describing age-dependent variation in aedeagal morphology in *A. uliginosus*, we adopt an integrative approach, evaluating internal morphological features, developmental characteristics, and COI-sequences to revisit the taxonomic relationship and status of *A. uliginosus* and *A. lotti*.

## Material and methods

### Morphological studies

As the initial step of the morphological analyses, we examined pinned dry specimens and ethanol-preserved specimens obtained from previous studies, from various regions including Germany (Baden-Württemberg, Bavaria, Rhineland-Palatinate, Saxony, Schleswig-Holstein), Austria, Hungary, and Southern Poland. The material was sourced from collections maintained by Konrad Dettner (Bayreuth, Germany), the Senckenberg Deutsches Entomologisches Institut (Müncheberg, Germany), and the Department of Hydrobiology at the University of Pécs (Pécs, Hungary).

Male specimens, regardless of their previous identification as *A. lotti* or *A. uliginosus* [hereinafter *A. uliginosus* s.l. (sensu lato)], were examined for their primary external sexual organs. In cases of 29 male specimens, internal male sexual organs were dissected and analysed in detail. Careful dissection was necessary to obtain intact phallobases, often involving the removal of tightly adhering basal parts of parameres, which were prone to damage during dissection. Age determination of freshly alcohol-preserved specimens from Hungary was conducted. The body weights of ethanol-preserved specimens from Hungary were measured one hour after deposition on filter paper using a Professional Digital Mini Scale, TL-series (1 mg – 20 g; Shenzhen Union Technology, China). Additionally, a few data on ectadenia were recorded from freshly collected specimens from Bavaria.

In the subsequent step, specimens were captured and kept alive from saline habitats of the Kardoskúti-puszta plain in Southeastern Hungary (42 specimens) on May 1, 2023. The living Hungarian specimens were immediately sent to Germany by express mail using fishing bait boxes filled with heavily moistened filter papers. After arrival, the specimens were frozen, and fresh weights were determined 30 minutes after defrosting. Sixteen male specimens were selected for detailed morphological analyses. Soft translucent chitinous structures at phallobases were dyed respectively darkened using a 1% aqueous or 1% ethanolic solution of pyrogallol (Seifert, 1995) for microscopic investigation. Darkening was controlled after 1 hour to 2 days to achieve optimal results.

To evaluate aedeagal morphology, measurements were taken on all available individuals, including the length of the penis from apex to distal corner of the basal part (a on Fig. 1A), the lengths of the penis base from the distal corner of the basal part to the basal part defined by the imaginary extension of the outer curvature of the penis on a line defined by a (b on Fig. 1A), the length of the sclerotized phallobase from the imaginary extension of the outer curvature of the penis to the basal sclerotized base of the phallobase in line with a + b (c on Fig. 1A), and the thickness of the non-sclerotized soft and translucent membrane of the phallobase (d on Fig. 1A). Basal pieces of both parameres and tegmenites (=tegmina) were also observed.

**Figure 1.**
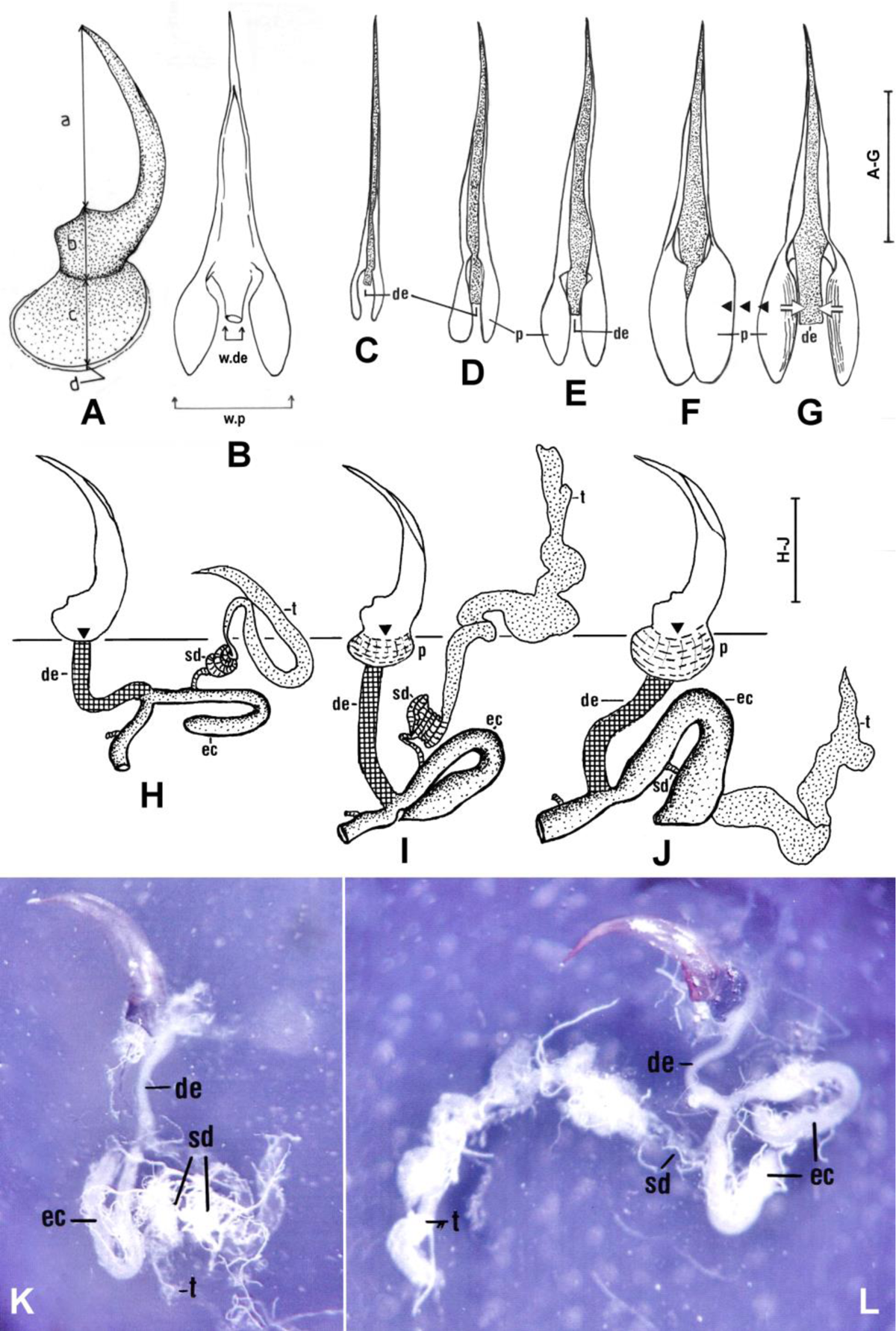
Genital morphology and its changes from young and elder specimens of *Agabus uliginosus* sensu lato. **A-B**: Morphology and aedeagus of an elder specimen in right side lateral (A) and dorsal (B) view, with measured lengths parameters a-d. **C-G:** Dorsal views of aedeagi from different age classes of *A. uliginosus* s.l.; C corresponds to *A. lotti*, while D-G to *A. uliginosus* s.str. with increasing age; C-E: Hungarian population; F-G: old male specimen from Rudna, Mühlgast, Poland; F and G show the same aedeagus isolated from a specimen preserved in ethanol, then dried for 5 minutes (F), and for 15 minutes or more (G). Black arrows on G symbolize movements of phallobases. **H-L:** Side view of aedeagi and morphology of internal genital organs of an immature (H, K, #H04), an elder (I, #H27) and an old (J, L, #H30) specimen of *A. uliginosus* s.l. from the Hungarian population. H and K correspond to *A. lotti*, J and L to *A. uliginosus* s.str, while I shows a young specimen of *A. uliginosus* s. str. as a transition between them. Abbreviations: de: ductus ejaculatorius (checkered in H-J), ec: ectadenies (dotted in H-J), sd: spermatic duct (cross-striped in H-J), p: phallobase, t: testes (dotted in H-J), w.p: width of phallobase, w.de: width of ductus ejaculatorius. The triangles and the interrupted horizontal line in H-J illustrate the origin of the phallobase as indicated by b in A. In K (#H04) and L (#H30) anterior margin and origin of ductus ejaculatorius is marked by arrows. Scales: 1 mm, different scales refer to A-G and H-J. For K and L, 20-fold magnification was used.

Out of the 45 male specimens of *A. uliginosus* s.l. examined in detail, fresh weight data were measured for 26 individuals (10 ethanol-preserved and 16 frozen-preserved, all from the Hungarian populations), while the most critical morphological parameters of the genitalia were recorded in 44 cases. All the beetles and genitals were examined, dissected, and measured using a ZEISS binocular and a ZEISS microscope (Standard 16). Phase-contrast photographs were captured using an OLYMPUS BH2-RFCA microscope. All morphometric measurements and visual inspection results were compiled in Table S1.

### Phenological records

To gain insight into the temporal distribution of records for both species, we conducted a comprehensive search for records with at least the month of collection. We utilized a variety of sources, including academic papers, specifically for *A. lotti* records: Turner et al., (2015), Boda, Móra & Csabai (2019, 2023), and Rocci, Terzani & Mascagni (2021). We accessed data from GBIF database (for specific dataset citations see Table S2), from verified public database of “Izeltlabuak.hu” citizen science initiative (http://www.izeltlabuak.hu) for both species, and obtained collection records from the Collection of Konrad Dettner (Bayreuth, Germany), the Senckenberg Deutsches Entomologisches Institut (Müncheberg, Germany), and the Collection of the Department of Hydrobiology of the University of Pécs (DHUP) for unpublished data on both species (Table S1).

### Molecular studies

#### DNA barcode amplification and sequencing

In cases of the 16 Hungarian male specimens that were maintained alive, subsequently frozen, weighed, and subjected to morphological analysis, we removed one hind leg from each specimen and preserved them in 96% ethanol, thus enabling us to conduct molecular studies (DNA barcode analysis). Samples were processed for sequencing at the Department of the Invertebrate Zoology and Hydrobiology, University of Lodz, Poland, by Zsolt Kovács and Tomasz Rewicz. DNA was extracted from a leg segment of the specimens using the Chelex procedure (Casquet, Thebaud & Gillespie, 2012). A 650 bp fragment of COI was amplified using primers LCO1490-JJ and HCO2198-JJ (Astrin & Stüben, 2008), under the following PCR conditions: initial denaturing for 60 s at 94°C, followed by five cycles of 30 s at 94 °C, 90 s at 45 °C, 60 s at 72 °C, 35 cycles of 30 s at 94°C, 90 s at 51 °C, 60 s at 72 °C, with a final 5 min extension at 72 °C (Hou, Fu & Li, 2007). PCR was conducted in 12 μL volumes, containing 1,1 μL of DNA template, 1 μL of each primer, 6 μL of DreamTaq PCR Master Mix (Thermo Scientific) and 2,9 μL of nuclease-free water. Then 2 μL of each reaction product was checked with 1% agarose gel electrophoresis to confirm amplification. 5 µL of PCR products were purified with Exonuclease I (2 U, EURx) and alkaline phosphatase Fast Polar-BAP (1 U, EURx), according to the manufacturer’s instructions. One-way Sanger sequencing was outsourced to Macrogen Europe (Amsterdam, the Netherlands). The sequences were edited, trimmed of primers, and aligned using Geneious 11.1.5 software package (Kearse et al., 2012).

#### DNA data assembly and barcode analyses

We used the BLAST (Altschul et al., 1990) searches to identify all the new sequences to confirm their identity based on already published sequences. All obtained COI sequences were deposited in GenBank (PP464831-PP464847; Table S3). Additionally, the DNA sequences were compared with 17 publicly available *A. uliginosus* s.l. and 1 *A. margareti* sequences retrieved from the public repository (Pentinsaari, Hebert & Mutanen, 2014, Hendrich et al., 2015, Rulik et al., 2017, Linard et al., 2018) of the Barcode of Life Data Systems (BOLD; Ratnasingham & Hebert, 2007), and altogether deposited in a separate DS-AGHUZSK dataset, where all the relevant metadata and sequence trace files will be publicly available upon publication (permanent link: http://www.boldsystems.org/index.php/Public_SearchTerms?query=DS-AGHUZSK, DOI after acceptance). Intra- and interspecific genetic distances were calculated based on the Kimura 2-parameter model (K2P; Kimura, 1980), using the analytical tools of the BOLD workbench (Distance Summary, and Cluster Sequences). A phenogram (see Table S3 for details) was constructed in MEGA X (Kumar et al., 2018) with the neighbor-joining method (Saitou & Nei, 1987), based on the K2P distance (Kimura, 1980), with a bootstrap test (1000 replicates). We estimated the genetic diversity, i.e., the haplotype diversity (h) and nucleotide diversity (*π*) (Nei, 1987), using the DnaSP v5 software (Librado & Rozas, 2009). The relationships within *A. uliginosus* were displayed through a median-joining network using PopART (Leigh & Bryant, 2015).

#### Species delimitation methods

We used three distinct species delimitation methods harboring two different approaches. Initially, we used two distance-based methods. First, we employed BINs (Ratnasingham & Hebert, 2013) integrated into BOLD, wherein COI DNA sequences, whether newly submitted or pre-existing, are organized and grouped into separate clusters according to their genetic distances. Secondly, we used the ASAP procedure, developed by Puillandre, Brouillet & Achaz (2021), which is a hierarchical clustering algorithm designed explicitly for species partitioning, leveraging pairwise distance distribution. The ASAP analysis was run with the iTaxoTools v.0.1 software (Vences et al., 2021). We applied the Bayesian implementation of the multi-rate PTP (mPTP - https://mcmc-mptp.h-its.org/mcmc/, Kapli et al., 2017) for the phylogeny-based delimitation method. This method incorporates MCMC sampling, offering a rapid and comprehensive evaluation of the deduced delimitation. We used Maximum Likelihood (ML) trees calculated with MEGA X (Kumar et al., 2018) as input for analysis.

## Results

### Morphological studies

Among the 45 male specimens of *Agabus uliginosus* s.l. subjected to detailed examination, a range of developmental stages were observed, including freshly hatched individuals with soft cuticles, very young, young, and more mature specimens (Table S1). Analysis of the genitalia revealed a correlation between age and the increasing lengths of phallobases (p), testes (t), and ectadenies (ec), which became progressively longer and thicker (Fig. 1H-J). The measurements (b+c) were found to be positively correlated with the lengths of ectadenies (Fig. 2B) and the fresh weights of the specimens (Fig. 2A), both in ethanol-preserved and frozen material, indicating that fresh weight roughly reflects the gradual enlargement of the phallobase. Additionally, the widths of phallobases in dorsal view (see Fig. 1C-G) significantly increased with age, along with the widths of ducti ejaculatori (Fig. 2C). Moreover, total lengths of aedeagi also showed an increase (Fig. 1C-J). This significant swelling process (Fig. 1C-H), observed from young to elder specimens, is noteworthy because it is unusual among adephagous beetles, as diameters in dorsal views of phallobases and whole aedeagi typically represent species-specific values. Consequently, changes in diameters of aedeagi in dorsal views, as well as varying shapes of the entire aedeagi ranging from straight to curved and even asymmetrical organs, were observed (Fig. 1C-H). Furthermore, an old, freshly dissected specimen (Fig. 1G) exhibited a different shape compared to the same specimen after drying (Fig. 1H).

**Figure 2.**
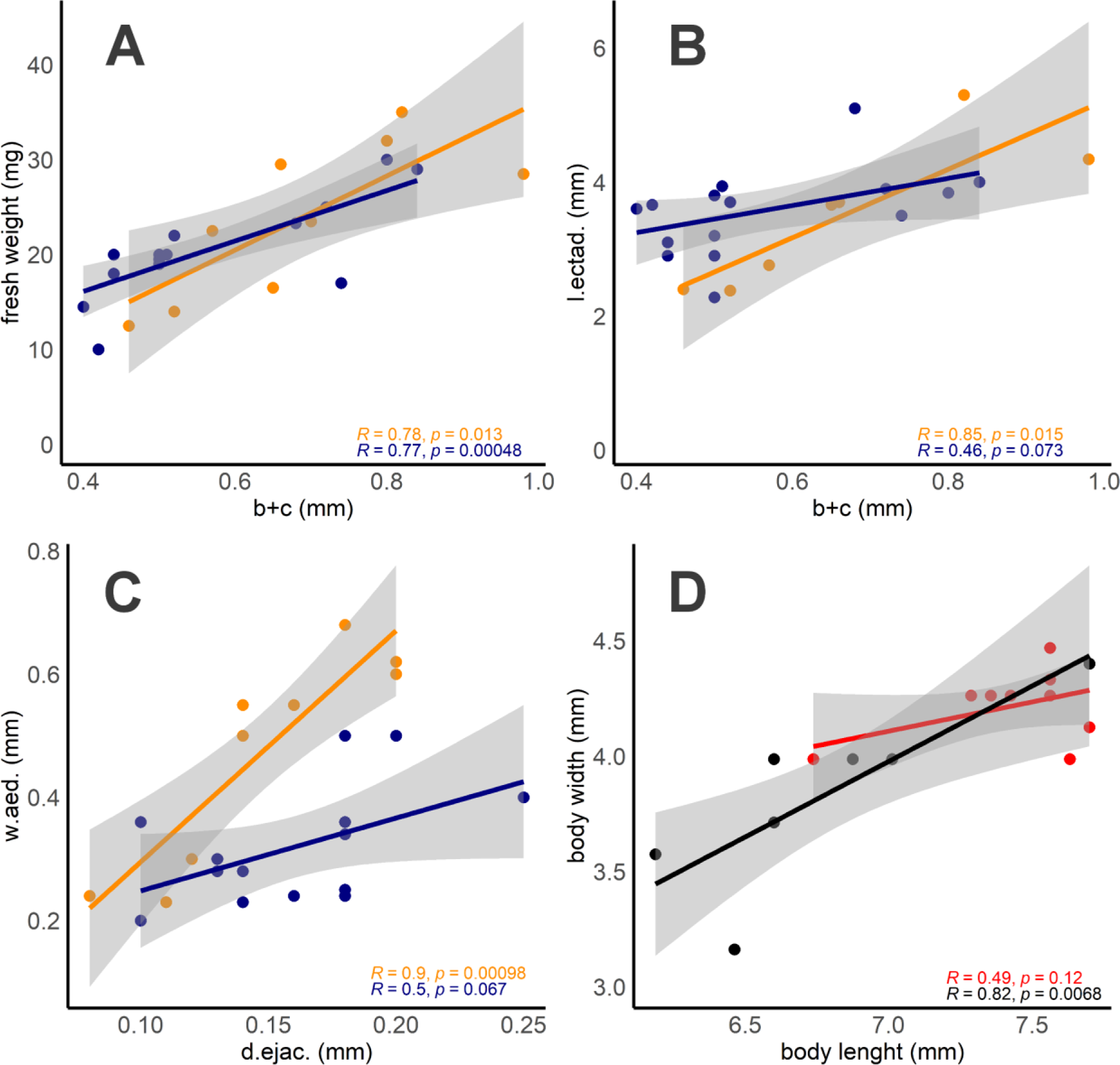
**A, B:** Fresh weights (A) and length of accessory glands (=ectadenies, l.ectad. here and ec in Fig. 1H-J) (B) as function of the size of the phallobase (b+c here and in Fig. 1A). **C:** Widths of the base of the aedeagus in dorsal view (w.aed.) compared with diameters of ducti ejaculatori (d.ejac.). A-C measurements were done in ethanol-preserved (orange) and frozen (blue) male specimens of *A. uliginosus* sensu lato from Hungary. **D:** Comparison of the body length and width of the males showing the differences and the variance in body shapes in *A. uliginosus* sensu stricto (red) and *A. lotti* (black) types.

Significant correlations were observed in *A. lotti* types, while marginally significant correlation can be seen in *A. uliginosus* s.str when comparing body length and width (Fig. 2D). However, the *A. uliginosus* s.str. specimens exhibited lower and less variable body lengths, with relatively constant widths. Conversely, in *A. lotti* type, both body length and width showed higher variability. We did not observe significant differences in body ratio (length/width) between *A. uliginosus* s.str. (mean=1.75, N=11) and *A lotti* types (mean=1.77, N=9) (Mann-Whitney U test, U=36.5, z=0.95, p=0.34).

Another distinctive characteristic observed in male representatives of the *A. uliginosus*-group is the soft and translucent external border of the phallobase (d in Fig. 1A), which is not sclerotized (Fig. 3A-C; arrows). Under higher magnification (Fig. 3C), radially arranged filaments were visible, indicating potential ongoing sclerotization processes along these structures. Older males, as shown in Fig. 3D, exhibited nearly complete sclerotization of section d, which is greatly reduced. Additionally, the sclerotized phallobase displayed both radial and circular filaments (Fig. 3D). The widths of these non-sclerotized phallobase borders (d) in elder and old male specimens varied distinctly. This unique morphological detail, optimally visible after artificial darkening with reagents, may suggest an extension of the phallobase even in old males, potentially indicating multiple gonadal activities over several months or even years. Consistent with these findings, both basal parts of parameres and tegmina broadened (tegmina also lengthened) and became increasingly sclerotized with age from youngest to the oldest, as shown on Fig. 4A-D, respectively.

**Figure 3.**
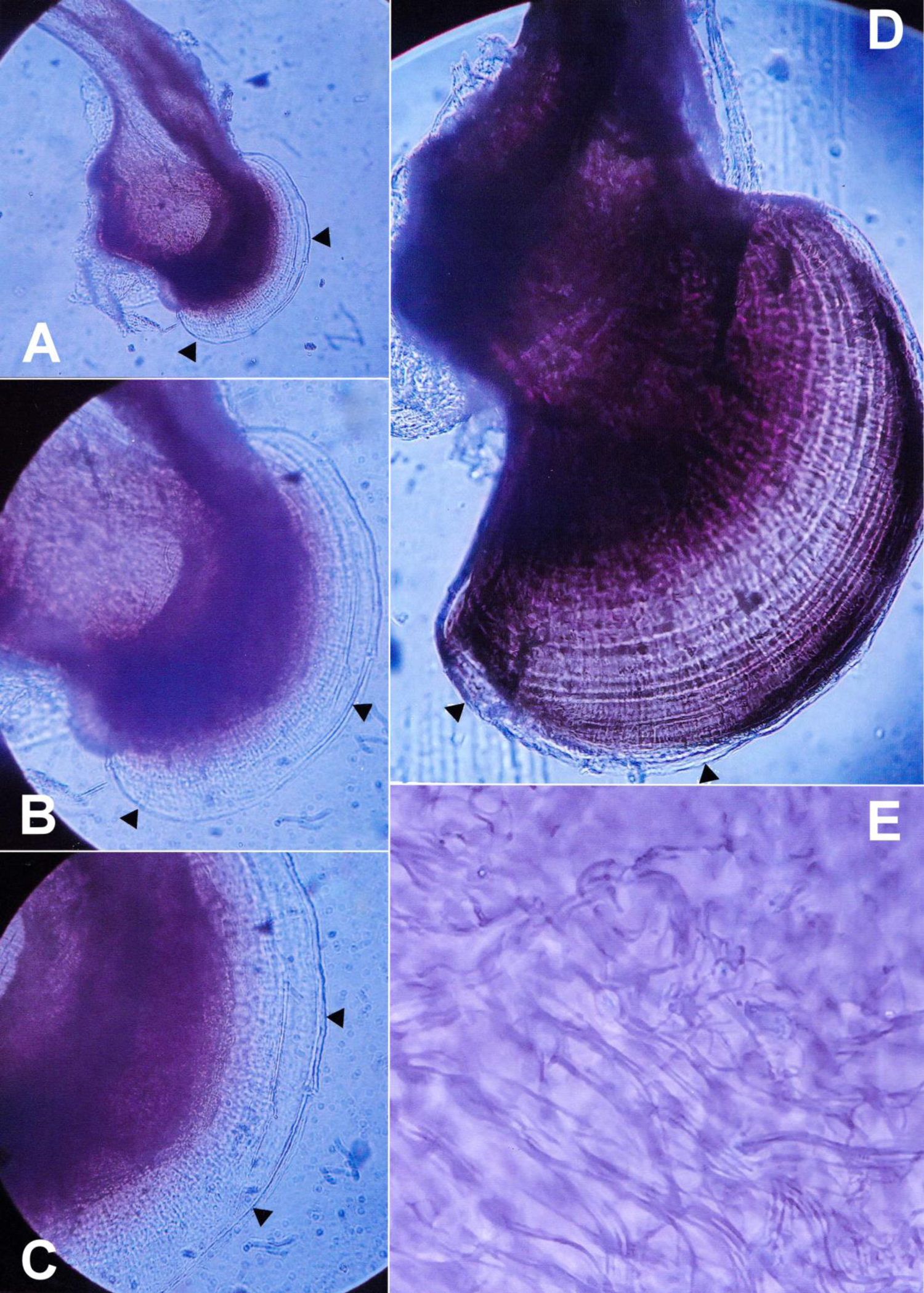
**A-D:** Translucent peripheral area of phallobase (d in Fig. 1A) of male *A. uliginosus* s.l. A-C: #G06 specimen from Crailsheim (Germany). D: Nearly completely sclerotized basal area of aedeagus of an old male *A. uliginosus* sensu stricto #P01 specimen from Rudna, Mühlgast, Poland with both radial and circular filaments. Arrows indicate the outer margin of d in Fig. 1A, radially arranged filaments between c and d are clearly visible, especially on C. **E:** Spermatozoans from spermatic duct of elder male #H27 from Hungary (see also Fig. 1I & 4C). Magnifications: A: 32-fold; B: 100-fold; C: 200-fold; D: 100-fold; E: 400-fold.

**Figure 4.**
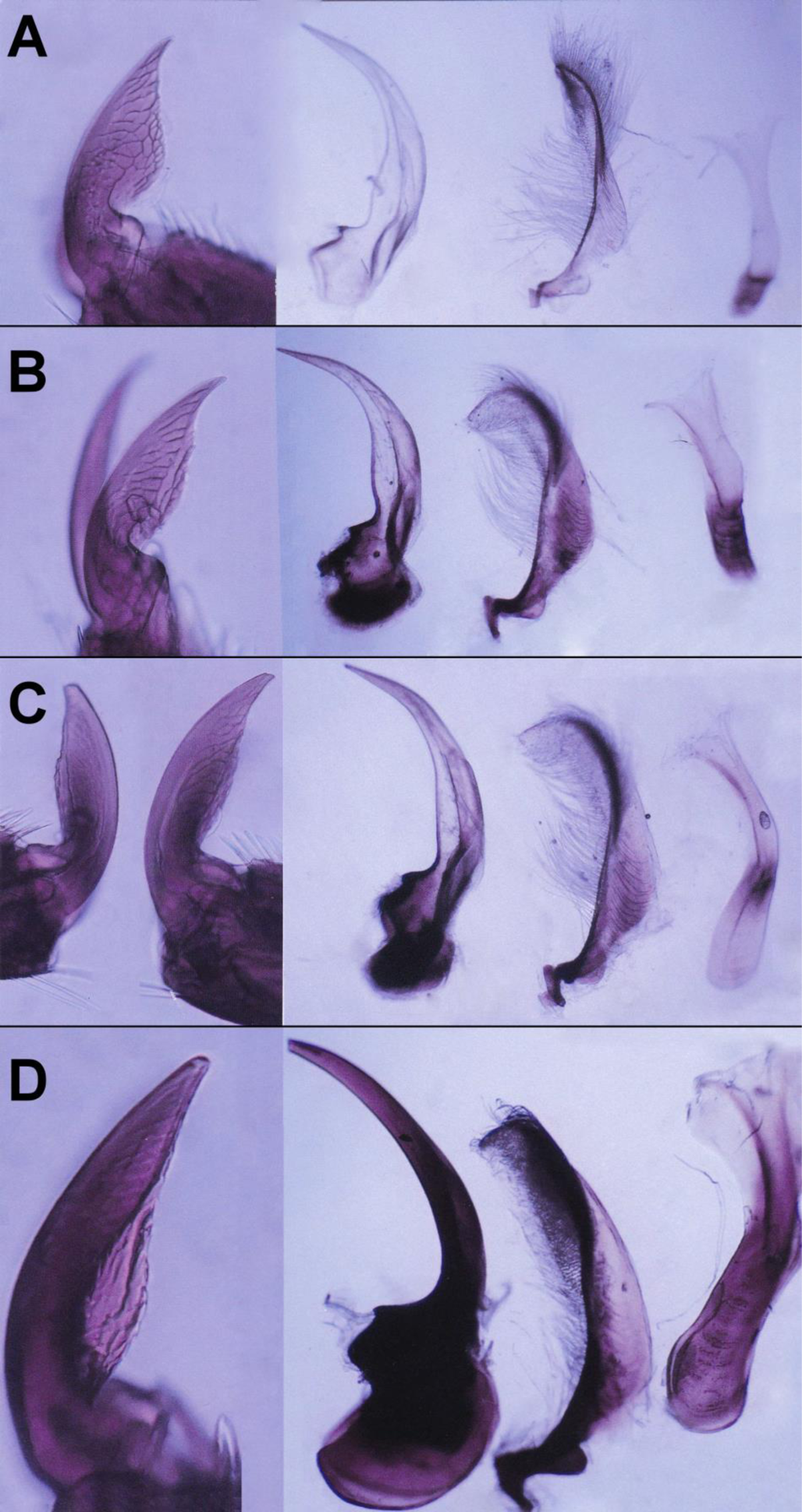
Morphological characteristics ((from left to right in all panels: foreclaws, aedeagi, parameres, tegmina) in *Agabus uliginosus* s.l. males of different age categories. A: #H19, B: #H11, C: #H27 specimens, all from Kardoskút, Hungary, D: #G02 from Salemer Moor, Ratzeburg, Germany. A: represent *A. lotti* type, B-D represent *A. uliginosus* s. str. type.

To sum up, during imaginal life, respective imaginal age classes of *A. uliginosus* s.l., the sclerotization of area c gradually increased, phallobases enlarged, parameres and tegmina lengthened, foreclaws became well-wore from immature specimens (see Fig. 1H, 3A-C, 4A-B) to elder and highly aged adults (Fig. 1I-J, 3D, 4C-D). Hereby, lengths below 0.6 mm (b + c) in immature specimens were tentatively associated with *A. lotti* (e.g., Fig. 1H & 1K). Conversely, males with b+c greater than 0.6 mm are interpreted as *A. uliginosus* s.str. (see Fig. 3, Fig. 1J & 1L). Based on genital morphology and size proportions, about half of the examined males (20 specimens) exhibited characteristics typical of *A. uliginosus* s.str. type, 21 specimens were of *A. lotti* type, while three individuals showed completely transitional characteristics and values.

### Phenology records

Altogether, we gathered more than 1600 dated records for *A. uliginosus* s.l. and 105 for *A. lotti* specifically. The majority of records for both species are concentrated in the months from March to June, with a notable peak in May. However, while data for *A. uliginosus* s.l. are spread across all seasons, with over 300 records from August to February, records for *A. lotti* are limited to the period from March to July (Table S4). Notably, only two records for *A. lotti* are available in July, one from Austria and one from Slovakia, likely originating from higher elevation sites.

### Molecular analyses

We successfully sequenced 16 specimens of *A. uliginosus* s.l. from Hungary. Comparison with all available *A. uliginosus* s.l. sequences in BOLD, revealed a relative homogeneity of the sequences regardless of whether our newly sequenced specimens morphologically classified to the *A. uliginosus* s.str. or *A. lotti* type.

The Neighbour-Joining tree (Fig. 5) and Median-Joining network (MJ network) (Fig. 6) clearly demonstrate that the newly sequenced individuals classified as ‘*A lotti*’ or *A. uliginosus* s.str are not distinct from each other or from the publicly available sequences in the BOLD database. While more ‘*A. lotti*’ sequences are located on the side branches of the MJ network, the two types remain intermixed throughout the entire network, with their haplotypes showing complete overlap (Fig. 6B).

**Figure 5.**
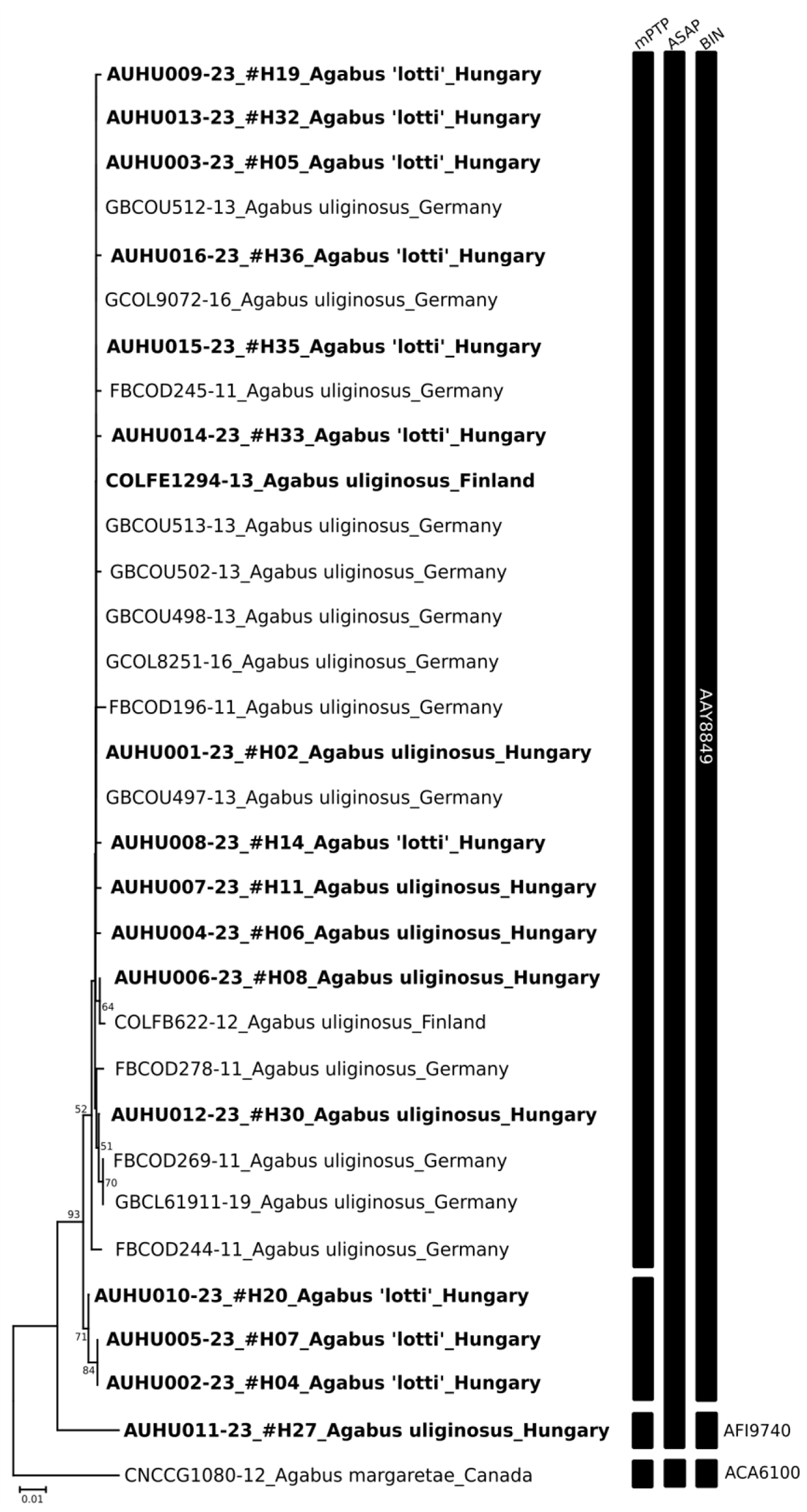
Neighbor-Joining tree based on COI K2P pairwise distances for sequences of *Agabus uliginosus* s.l. and *Agabus margareti* (name written as in BOLD) used as an outgroup. Names written in bold have genitals checked for morphological discrimination between ‘lotti’ and ‘uliginosus s.str.’ types. Each sequence name consists of parts separated by a lower dash: (1) BOLD Process ID, and for the Hungarian specimens the specimen codes used in morphological analyses are also given, (2) species (types), and (3) country of origin. The results of the three species delimitation methods are indicated by vertical bars (BIN numbers given). Only bootstrap supports above 50% are shown.

**Figure 6.**
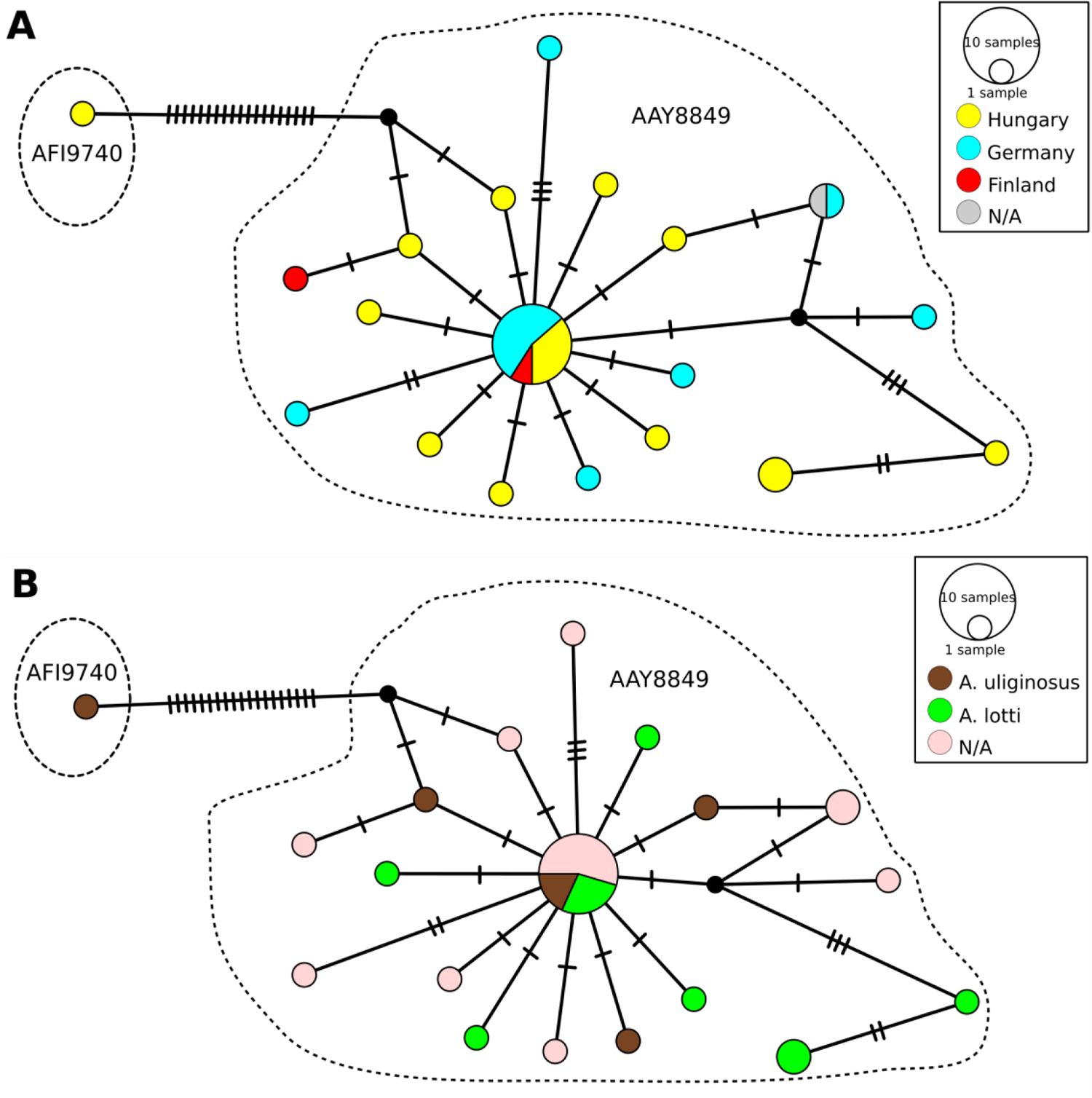
Median-Joining network showing the relationships among the haplotypes of *Agabus uliginosus* s.l. in the context of: A) country of origin, B) morphotypes. Colors indicate individuals representing different countries of origin or morphological type. Each bar represents one substitution, whereas small black dots indicate undetected/extinct intermediate haplotype states. The sizes of the circles are proportional to the frequencies of haplotypes (see open circles with numbers). Dashed lines encircle separate BINs.

In our sample from a single Hungarian population, we identified 12 haplotypes, with an additional seven haplotypes from public sequences in BOLD, resulting in a total of 19 known haplotypes and two intermediate haplotypes in the entire network (Fig. 6). From Germany, seven, and from Finland, two haplotypes were identified. The central haplotype, comprising 10 sequences, includes individuals from all three countries (Fig. 6A) and both types (Fig. 6B), i.e. the Hungarian individuals classified as both *’A. lotti*’ and *A. uliginosus* s.str., sequences that are 100% identical to each other and to sequences from Germany and Finland. Apart from the central haplotype, two haplotypes are represented by two copies, while the others are represented by only one.

Species delimitation methods yield consistent classifications for most sequences, with only a discrepancy observed in the case of three Hungarian sequences classified as ‘lotti’ type and one as uliginosus s.str type. The most conservative method (ASAP) assigns all sequences to a single Molecular Operational Taxonomic Unit (MOTU). For delimitation based on BINs, all but one new sequence aligned to the BIN BOLD:AAY8849, which already contained all 17 previously known *A. uliginosus* s.l. sequences. Within this relatively compact and homogeneous BIN, the average distance was 0.38%, and the maximum distance was 1.52%. However, the specimen #H27 was grouped separately and formed a singleton BIN (BOLD:AFI9740), which became the nearest neighbour of the *A. uliginosus* BIN, with a distance of 3.7%. In comparison, the only species of the species group whose sequence was available in the BOLD (*A. margareti,* BOLD:ACA6100 singleton BIN) showed a distance of 6.9% to its nearest neighbour, which, like that of the new #H27 BIN, was also the ‘original’ *A. uliginosus* BIN. Adding specimen #27, so including both *A. uliginosus* BINs, the average within-species distance increased to 0.72%, while the maximum within-species distance was 4.57% (Table S5.).

## Discussion

### Morphological characterization of A. uliginosus sensu lato, including A. lotti

Among central European *Agabus* species, *A. uliginosus* is characterized by metacoxal lines fully reaching the hind margin of the metasternum and broad metasternal wings, according to Schaeflein (1971) and Nilsson & Holmen (1995). Additionally, the anterior pronotal puncture lines are continuous and not distinctly interrupted. The meshes of elytral sculpture are typically small, smaller than elytral punctures, as noted by Ganglbauer (1892). Balfour-Browne (1950) provided a concise key to differentiate *A. uliginosus* from similar species such as *A. sturmii* (Gyllenhal, 1808), *A. unguicularis* (Thomson, 1867), *A. labiatus* (Brahm, 1790), and *A. congener* (Thunberg, 1794). In *A. sturmii*, the body length is about 8 mm, and anterior male foreclaws lack teeth. In the other mentioned species, including *A. uliginosus*, body lengths are maximally 7.6 mm. Metasternal wings are distinctly narrow in *A. unguicularis* and *A. labiatus*, whereas they are broad and triangular in *A. congener* and *A. uliginosus*. A characteristic tooth is present on the anterior male claws of foretarsi in *A. uliginosus* but absent in *A. congener*. Furthermore, the complete prosternal ridge is roof-like in *A. uliginosus*, while it fades out to an arched process in *A. congener*. Additionally, the metatibial hind spine is as long as the first metatarsal segment in *A. uliginosus*, but longer in *A. congener*. Characteristically, *A. uliginosus* exhibits a strongly convex dorsal surface and very broad pronotal lateral margins. Nilsson & Petrov (2005) separated *A. uralensis* from *A. uliginosus*, as a vicariant species in the western Palaearctic, based on its smaller body size and shorter, evenly tapering penis.

As several authors before, recently Turner et al. (2015) noted high variance in several external morphological characteristics of *A. uliginosus*, including microsculpture and coloration. Stephens (1828) described stronger metallic specimens from England, while Queney (2002) observed a rufous form from France. Moreover, *A. uliginosus,* uniquely in the subfamily Agabinae, shows inter- and intrasexual dimorphism within part of its distribution area (Bilton et al., 2016).

### Biology and population structure of A. uliginosus

The life cycle of *A. uliginosus* is characterized as an univoltine spring breeder with summer larvae and overwintering adults (Nilsson, 1986). Galewski (1968, 1980) observed second-stage larvae from early May to late July in Poland, followed by third-stage larvae from late April to mid-September. He noted the presence of mostly third-stage larvae but also pupae and immature adults during May, suggesting a prolonged oviposition period. Nilsson (1986) suggested the possibility of an extended oviposition period as well. Similar bionomical data were reported by Balfour-Browne (1950). In contrast, Braasch (1989) proposed that *A. uliginosus* has two oviposition periods: the first from March to April and the second from July to August in Northeastern Germany. He demonstrated two peaks of adult beetles, one from January to March after adult hibernation and another from the end of May to October. Two peaks were also observed in both second- and third-stage larvae, with the first peak from April to the beginning of May and the second peak in July. Braasch (1989) also noted that *A. uliginosus* larvae are typically found in temporary waters, whereas adults inhabit permanent water bodies. Spitzenberg (2021) reported a phenological maximum for *A. uliginosus* during May in Saxony-Anhalt and observed avoidance of higher elevations. He also described the ecological preferences of the species for eutrophic ponds, backwaters, ditches, canals, salt lakes, and salt marshes in this area. Additionally, Spitzenberg (2021) reported flight activity at light, a behaviour supported by Kehl and Dettner (2007), who associated this species with category 2a, indicating the presence or absence of flight muscles depending on individuals.

Summarizing available records from Central Europe, we observe a peak in records during March-April-May for both species, consistent with the phenological observations mentioned above. Additionally, the exclusive presence of specific records of *A. lotti* during these months supports the hypothesis that this species may actually represent young individuals of *A. uliginosus* returning to the water shortly after hatching, potentially contributing to the surge in density, and consequently in number of records. However, it is essential to consider that climatic variations and differences in altitude make objective phenological comparisons challenging. For instance, March marks the beginning of the breeding season in southern and lowland areas, whereas reproduction may commence later, typically in May, in higher elevations. Additionally, data on A. ‘lotti’ have been documented in mountainous regions of Italy, Austria, and Slovakia.

As demonstrated by Classen and Dettner (1983) and Dettner, Hübner & Classen (1986), age structures within dytiscid populations can exhibit significant fluctuations from month to month, over the course of one or several years, although the exact lifespan of many Dytiscid species, including *A. uliginosus*, remains unclear. In species such as *Agabus bipustulatus* (Linnaeus, 1767), *A. paludosus* (Fabricius, 1801), or *Platambus maculatus* (Linnaeus, 1758), populations typically consist of varying numbers of individuals across different age classes at any given time, and these proportions are constantly changing. These age classes are delineated based on the developmental status of internal gonads, which can only be reliably assessed in freshly killed or recently ethanol-preserved specimens. Therefore, any additional records or notes accompanying regular distribution data regarding the first occurrences of larval stages or immature (soft, unsclerotized) adults or information on the status of the gonads provide crucial information for determining the life-cycle type of each species, in accordance with Nilsson (1986).

### Age dependency of male sexual organs of A. uliginosus

Within the *uliginosus*-group, and likely in the *punctulatus*-group as well, Nilsson (2000) suggested that during the imaginal stage, the degree of sclerotization and enlargement of the phallobase of the aedeagus might increase. Nilsson & Petrov (2005), after studying a larger material of *A. uliginosus* and *A. uralensis*, concluded that identification might be problematic if the phallobase did not enlarge since the development of the basal apodeme occurs later than that of the rest of the penis. This is unusual because sclerotized aedeagi of male insects typically represent morphologically stable, constant, and highly useful diagnostic structures, allowing species differentiation in most insect groups. The lock-and-key mechanism (Eberhard, 1985) posits that the morphologically species-specific external genitalia of older specimens correspond tightly between both sexes. However, the aedeagal morphology of freshly hatched male beetles has not been systematically analysed before and has not been compared with appropriate structures of older specimens, including the *A. uliginosus*-group.

As reported by Classen and Dettner (1983) in *A. bipustulatus* and *A. paludosus*, it was expected, and now demonstrated for the first time in *A. uliginosus* s.l. (Fig. 1H-J), that the lengths and diameters of ectadenies gradually increase from one age class to another.

Additionally, and in line with Dettner, Hübner & Classen (1986), the fresh weights also positively correlated with the lengths of “b+c” (Fig. 2A, see also Fig. 1A). At the same time, we clearly demonstrated that aedeagal morphology gradually changes with age: the phallobase enlarges, its basal part becomes increasingly sclerotized, the parameres and tegmina lengthen from immature, young specimens (see Fig. 1H, 3A-C, 4A-B) to older and highly aged adults (Fig. 1I-J, 3D, 4C-D). During this process, younger adults resemble *A. lotti* more, while older and fully sclerotized adults resemble *A. uliginosus* s.str. more. Within Dytiscidae, beside the *Agabus uliginosus*-group, Nilsson and Petrov (2005) mentioned a comparable case in *Hygrotus* (*Leptolambus*) *tumidiventris* (Fall, 1919), with a normal aedeagus figured by Larson (1975).

While they are not explicitly sexual organs, the foreclaws of males also play a role in the mating process, thus becoming worn out over age classes, with their apex becoming blunted, deformed, and eventually breaking off. We hypothesize that the fixing and adhering of male foreclaws to the pronoti of females during one or more extended copulations, particularly with increasing age, may damage both the thin and sensitive arched apodemes situated at the base as well as the tips of the foreclaws. Our results also underscore the urgent need for comparing aedeagal morphologies of young teneral males with those of adult specimens, not only in Dytiscidae but also in other taxa.

This is particularly crucial for older descriptions where figures of aedeagi were provided in dorsal views. When large amounts of viscous ectadenial materials, especially proteins and mucosubstances, as observed in Carabidae (see Krüger et al., 2014; Schubert et al., 2017), pass through aedeagi, these structures, if not completely sclerotized, may swell enormously. Depending on the amounts and consistency of ectadenial secretions, a certain flexibility of aedeagal structures is essential.

### Molecular diversity and MOTU delimitation

DNA barcoding is already a well-established and powerful tool for molecular identification of the species (Grant et al., 2021; Antil et al., 2023). Beetles are among the flagship groups of invertebrates for which barcode reference libraries are built (Pentinsaari et al., 2014; Hendrich et al., 2015). Still, even in Europe, progress reached above 50% of known species for freshwater aquatic beetles (Csabai et al., 2023).

Species delimitation methods revealed two (ASAP), three (BIN), and four (mPTP) MOTUs, these also considered morphological features and the accepted approach can be considered as a rough proxy of species. We should consider the methods’ limitations and interpretation of the obtained result. The conservative approach of the ASAP method seems congruent with our morphological investigation (Copilaş-Ciocianu et al., 2022), and the mPTP approach tended to oversplit (Song et al., 2018, Goulpeau et al., 2022). While the phenomenon of bin-sharing may arise here (e.g., Raupach et al., 2018; Raupach et al., 2020; Schattanek-Wiesmair et al., 2024), we can rule out this possibility since in some cases, the sequences are 100% identical for individuals classified morphologically as different types. This holds even when comparing Hungarian *A. lotti* individuals with German and Finnish *A. uliginosus* s.str.

As a theoretical possibility, it could be argued that all previous COI sequences in BOLD may have originated from *A. lotti*. However, this can be clearly ruled out based on the attached photos of processed individuals in BOLD, especially with reference to the specimen with Process-ID COLFE1294-13, where the genitals (aedeagus and parameres) are visible, leaving no doubt that this old specimen belongs to *A. uliginosus* s.str.

The addition of the Hungarian sequences significantly increased the known haplotype diversity within the species. Large number of individuals are available from Germany, Finland, and Hungary in BOLD, and the Hungarian dataset stands out as the most heterogeneous. Only available molecular data from *A. lotti* are deposited in Venables’ thesis (2016). One specimen from Italy is closely related to *A. uliginous* from Germany. However, it is noteworthy that Venables utilized COI-3P sequences, and unfortunately, the sequence corresponding to *A. lotti* is not publicly accessible in repositories. Despite this limitation, the observed genetic divergence appears relatively modest. Given the geographical context (Italy), it is plausible to consider the possibility of intraspecific variation induced by spatial isolation, although definitive conclusions warrant further investigation.

The comparison of sequences confirmed with high confidence that individuals classified as *A. lotti* and *A. uliginosus* belong to the same species based on COI markers. Although it nicely supports our single-species hypothesis in the present case, the COI analysis alone is insufficient for a comprehensive revision of the species complex. It would be highly beneficial to conduct a comprehensive multi-marker study encompassing most of the distribution range of the species group, covering various regions of the Holarctic region, including all known species (currently 6 without *A. lotti*), and employing both molecular and morphological approaches integratively, including analysis of immature, young, and teneral individuals. Limited sampling size may also affect our findings.

### The case of the #H27 specimen

As shown, specimen #H27 (morphologically identified as *A. uliginosus s.str.*) displayed molecular deviations from all other specimens, irrespective of their morphological identification as *A. uliginosus* s.str. or *A. lotti*. Therefore, detailed comparisons were made of the aedeagi, parameres, tegmina, male foreclaws, and internal gonads of the molecularly analysed specimens. Young specimens were characterized by non-sclerotized, slightly coloured, translucent genital sclerites, small and lengthened tegmina, as well as thin and small bases of parameres (Fig. 4A right; #H19). In contrast, elder specimens exhibited thickened and more sclerotized tegmina, along with thickened and sclerotized bases of parameres (Fig. 4B right; #H11). Specimen #H27 (Fig. 4C right) was notably older compared to #H19 (Fig. 4A right), displaying typical features of *A. uliginosus* s.str., including an increased phallobase and parameres. However, #H27 had a lengthened tegmen that was only partially sclerotized in the center and not at the base. Exhibiting non-sclerotized internal organs, #H27 (c+d= 0.42 mm; d.ej.: 0.18 mm; ect.: 3.5 mm) appeared slightly younger than #H11 (c+d= 0.34 mm; d.ej.: 0.25 mm; ect.: 5.1 mm, see also Table S1).

Nevertheless, based on its sclerotized and broader phallobase, #H27 was conclusively older than #H11. Age determination was further supported by the exclusive presence of sperms in the spermatic ducts (sd) of #H27 (Fig. 3E), contrasting with their absence in #H11 and #H19. Although specimens from the Hungarian population were not very old, external genitalia were compared to distinctly older *A. uliginosus* specimens from other regions (despite the unavailability of internal genitalia). Two specimens (#G01, #G02) from Salemer Moor/Ratzeburg, Germany, both with c+d-values of 0.4 mm, large phallobases, and thick and sclerotized basal pieces of parameres, exhibited characteristic lengthened tegmina similar to those seen in #H27. This comparison further confirmed that Hungarian specimen #H27 represented an older specimen of *A. uliginosus* compared to #H11.

Furthermore, the larger asymmetrical foreclaw was compared among these three Hungarian and one German male specimens (Fig. 4A-D left). The youngest specimen, #H19, displayed a highly arched basal apodeme with a very sharp tip (Fig. 4A left). In the somewhat older specimen #H11, the basal apodeme of the sharp claw (only one leg was available) appeared slightly slender and frayed out (Fig. 4B left). Specimen #H27 was characterized by a worn-out basal apodeme and rounded or broken tips of the claws (Fig. 4C left), indicating that #H27 is the eldest specimen among the three Hungarian specimens depicted. The old specimen from Germany exhibited a male foreclaw with a rounded tip and a slightly frayed basal apodeme (Fig. 4D left).

The case of specimen #27 unequivocally indicates that while current knowledge and examination of northern populations have revealed a relative genetic homogeneity, parts of the distribution area not yet genetically characterized, particularly the southern and eastern areas, may harbour unexpected discoveries, potentially exhibiting a broader diversity of haplotypes and even revealing previously unknown hidden species. As shown by Bergsten et al. (2012), in a comprehensive study 250 individuals were needed to demonstrate 95% of the molecular variation of the widespread species *Agabus bipustulatus*, and around 70 for other species with more restricted distribution. More extensive sampling in the Pannonian Basin may fill the gaps regarding *A. uliginosus* group, as this region is proven to be a biodiversity hotspot for several aquatic invertebrate groups (Csapó et al., 2020; Sworobowicz et al., 2020). With limited sampling, we cannot exclude or confirm the phylogeographic effect of molecular separation of specimen #27. Multiple BINs within species are not unique (Raupach et al., 2020; Geiger et al., 2021).

### Conclusions and the taxonomic position of Agabus lotti

All our findings indicate that beetles identified as *Agabus lotti* represent young specimens of *A. uliginosus.* Therefore, the former should be regarded as a junior synonym of the latter. We observed and demonstrated a gradual morphological transition of aedeagal features from young specimens (*A. lotti*) to elder individuals (*A. uliginosus*), corresponding with age-dependent variations of internal sexual organs. Obviously, Turner et al. (2015) was unable to analyse both sclerotized and soft structures across a series of male specimens of different ages. However, they noted in their description of *A. lotti* that the endophallus varies morphologically depending on the maturity of the beetles and referenced remarks by Nilsson and Petrov (2005), who observed enlarged aedeagi in mature *A. uralensis* specimens and underdeveloped ones in young teneral specimens. Additionally, the fact that nearly all localities of *A. lotti* are found within the geographic range of *A. uliginosus* (with exceptions possibly in Central Italy) further challenges the species priority of *A. lotti*. Our molecular analyses further reinforced our hypothesis that *A. lotti* and *A. uliginosus* are the same species, as we observed only minimal differences among most specimens, with all but one individual classified into a BIN demonstrating high homogeneity. However, the genetically distinct nature of a single individual also underscores the need for further investigations in the species group. Particularly, with regards to the possible existing but never analysed southern populations, more significant differences may exist, or the possibility of discovering new, previously unknown species cannot be ruled out.

## Acknowledgements

KD is thankful to members of Entomological Society (Stuttgart), who supported him so much during many years, especially Hans-Ulrich Kostenbader (+, Stuttgart), Jürgen Frank (Waiblingen-Beinstein), and Eberhard Konzelmann (Ludwigsburg). Furthermore, he thanks Dr. Karl Wilhem Harde (+, Natural History Museum, Stuttgart) and Prof. Dr. Hinrich Rahmann (Zoology University of Stuttgart-Hohenheim, Hagen) for their enormous support. Collection material was gratefully lent from the collection of the Senckenberg Deutsches Entomologisches Institut (Müncheberg; Mrs. Mandy Schröter, Dr. Stephan Blank, and Dr. M. Simoes) and from Bernhard Moos (Auerbach). Prof. Dr. Sandra Staiger and PD Dr. Johannes Stökl (Chair of Evolutionary Animal Ecology, University of Bayreuth) gratefully allowed the use of their optical facilities. Authors are thankful to Łukasz Trębicki (University of Lodz) for assistance in the molecular laboratory. ZK was supported by ERASMUS 22/1/KA131/HED000057355/SMT-094 and CEEPUS F-2324-179374 grants. KD thanks Brigitte Dettner for participating and supporting him during various excursions. Authors used ChatGPT 3.5 for checking linguistic correctness and improvement of the readability of several paragraphs, but not a single sentence, text fragment, or independent thought was initiated or generated using large language models. Authors reviewed, revised and made decisions on the ChatGPT suggestions to their own liking and take ultimate responsibility for the content of this publication.

## SUPPORTING INFORMATION

Additional supporting information (5 tables) can be found online in the Supporting Information section at the end of this article.

